# Identification and characterization of moonlighting long non-coding RNAs based on RNA and protein interactome

**DOI:** 10.1101/261511

**Authors:** Lixin Cheng, Kwong-Sak Leung

## Abstract

Moonlighting proteins are a class of proteins having multiple distinct functions, which play essential roles in a variety of cellular and enzymatic functioning systems. Although there have long been calls for computational algorithms for the identification of moonlighting proteins, research on approaches to identify moonlighting long non-coding RNAs (lncRNAs) has never been undertaken. Here, we introduce a methodology, MoonFinder, for the identification of moonlighting lncRNAs. MoonFinder is a statistical algorithm identifying moonlighting lncRNAs without a priori knowledge through the integration of protein interactome, RNA-protein interactions, and functional annotation of proteins. We identify 155 moonlighting lncRNA candidates and uncover that they are a distinct class of lncRNAs characterized by specific sequence and cellular localization features. The non-coding genes that transcript moonlighting lncRNAs tend to have shorter but more exons and the moonlighting lncRNAs have a localization tendency of residing in the cytoplasmic compartment in comparison with the nuclear compartment. Moreover, moonlighting lncRNAs and moonlighting proteins are rather mutually exclusive in terms of both their direct interactions and interacting partners. Our results also shed light on how the moonlighting candidates and their interacting proteins implicated in the formation and development of cancers and other diseases.

## 1. Introduction

Protein moonlighting is a common phenomenon in nature involving a protein with a single polypeptide chain that can perform more than one independent cellular function (Boukouris et al, 2016; Monaghan & Whitmarsh, 2015). Enzymes, receptors, ion channels or chaperones are the typical form of moonlighting proteins (MPs). Enzyme is the most common form of moonlighting proteins whose primary function is enzymatic catalysis, but they are also in possession of additional roles such as signal transduction, transcriptional regulation, apoptosis, motility, and structural proteins (Jeffery, 2015). For example, *crystallins*, a class of well-studied moonlighting proteins, function as enzymes when expressed at low levels in many tissues, but are densely packed to form lenses when expressed at high levels in eye tissue (Piatigorsky et al, 1988; Piatigorsky & Wistow, 1989). The genes encoding *crystallins* need to sustain functions of both catalytic and transparency maintenance. Another example is glycolysis, an ancient universal metabolic pathway, in which a high percentage of proteins are moonlighting proteins (Boukouris et al, 2016; Sriram et al, 2005). Moreover, some proteins work on their moonlighting by being assembled to supramolecular, such as the ribosome, which usually composed of more than a hundred of proteins and RNAs. However, the studies of moonlighting merely concentrated on proteins and the genes coding them, yet the moonlighting of non-coding RNAs has not been investigated, despite the fact that ncRNAs have gained widespread attention due to their functional importance over the last decade (Chen, 2016b; Liao et al, 2011; Quinn & Chang, 2016; Zhou et al, 2017a).

Currently, the information of MPs, such as protein functions, cell localization, and primary structures, is scattered across a number of publications, since the MPs perform a variety of functions in different tissues and cell types. Some researchers have summarized the literature about MPs from different aspects of the functional diversity, such as regulation circuits or signaling pathways. The Jeffery lab constructed a manually curated database MoonProt, which consists of over 200 MPs that have been experimentally verified (Mani et al, 2015). The structures and function information about the MPs can aid researchers to understand how proteins function in a moonlighting manner and help in designing proteins with novel functions. Min *et al.* summarized the MPs from the perspective of a coupled intracellular signaling pathway (Min et al, 2016). Numerous proteins are localized in more than one compartment in cells and the aberrant translocation of proteins may cause cancer or other disorders. Hence, it is necessary to study the localization dynamic and trans localization activity of MPs. Monaghan *et al.* reviewed several MPs with dual mitochondrial and nuclear functions (Monaghan & Whitmarsh, 2015). It is pointed out that the nuclear moonlighting of mitochondrial proteins is part of a mitochondria-to-nucleus signaling pathway in cells. They also discussed various mechanisms commanding the dual localization of proteins and indicated that the nuclear moonlighters perform as a regulating loop to maintain homeostasis in mitochondria. Boukouris *et al.* summarized the moonlighting functions of metabolic enzymes in the nucleus (Boukouris et al, 2016). They proposed a new mechanistic connection between metabolic flux and differential expression of genes, which is implemented via nuclear translocation or moonlighting of nuclear metabolic enzymes. This mechanistic link aids cells in adapting a changing environment in normal and disease states, such as cancer, and thus has the potential to be explored for novel therapeutic target.

In parallel to the serendipitous findings of MPs through experiments, some computational approaches have been developed to predict MPs in recent years (Pritykin et al, 2015). Specifically, three algorithms were proposed for moonlighting protein identification, MoonGO (Chapple et al, 2015), MPFit (Khan & Kihara, 2016), and DextMP (Khan et al, 2017), executing statistic, machine learning, and text mining techniques, respectively. These studies investigated different aspects of MPs such as conserved sequence domains, structural disorders, protein interaction patterns, and network topology. MoonGO first identifies overlapping protein clusters in the human interactome (Chapple et al, 2015). Then, the clusters are annotated to the Gene Ontology (GO) terms of biological process. GO terms annotating more than half of a cluster’s proteins are assigned to the cluster. Each individual protein then inherits the annotations of its clusters in addition to its own. Finally, the proteins shared by dissimilar functional clusters are identified as MPs. MPFit uses a variety of protein features to address the diverse nature of MPs, including functional annotation, protein interactions, gene expression, phylogenetic profiles, genetic interactions, network-based graph properties, and protein structural properties (Khan & Kihara, 2016). In general, MPs are assigned in more clusters because they interact with proteins of diverse functions, so the number of clusters that a protein involved is used as an omics feature. For proteins that do not have an available record of certain features, an imputation step using random forest is used to predict the missing features. Eventually, these features are combined with machine learning classifiers to make moonlighting protein prediction. DextMP is a text mining algorithm consisting of four logical steps (Khan et al, 2017). First, it extracts textual information of proteins from literature and functional description in the UniProt database. Next, it constructs a k-dimensional feature vector from each text using three language models, *i.e.*, paragraph vector, Term Frequency-Inverse Document Frequency (TFIDF) in the bag-of-words category, and Latent Dirichlet Allocation (LDA) in the topic modeling category. Third, using four machine learning classifiers, a text is classified to either MP or non-MP based on the text features. Finally, the text predictions for each protein are separately summarized to predict which ones are MPs.

Long non-coding RNAs (lncRNAs) is a subclass of non-coding RNAs with little coding potential whose transcript consists of no less than 200 nucleotides. lncRNAs are implicated in a variety of biological processes through diverse functional mechanisms such as chromatin remodeling, chromatin interactions, and functioning as competing endogenous RNAs (Ferre et al, 2016). Specific expression patterns of lncRNA in cells correspond to certain cell development and disorder. Nuclear and cytoplasmic lncRNAs can regulate gene expression and function in multiple ways, e.g., 1) affecting mRNA translation directly, 2) interfering with protein post-translational modifications to disturb signal transduction, 3) functioning as decoys for miRNAs and proteins, 4) acting as miRNA sponges, 5) interacting proteins to enhancer regions, and 6) encoding small proteins and functional micro peptides, *etc*. (Cabili et al, 2015; Ferre et al, 2016; Quinn & Chang, 2016; Zhou et al, 2017b; Zhu et al, 2016). Many lncRNAs diversely reside in the nucleus and play an essential role as modulators for nuclear functions. Some others are translocated to the cytoplasm to enforce their regulatory roles. In some cases, these lncRNAs are implicated in an anterograde pathway bridging the nucleus and the mitochondria. Moreover, lncRNAs have a variety of subcellular localization patterns, which are not limited to specific nuclear and cytoplasm localization but also nonspecific localization in both the nucleus and cytoplasm (Barabasi & Oltvai, 2004; Buxbaum et al, 2015). For the lncRNAs localized in multiple compartments, the intercommunication can modulate the interaction pattern or expression abundance, e.g. regulating the lncRNA abundance in one compartment may influence the function of the other cell compartment. Also, inappropriate moonlighting may trigger genetic diseases (Abumrad & Lange, 2006; Espinosa-Cantu et al, 2015; Min et al, 2016). Hence, it is necessary to study the localization dynamic and expression activity of moonlighting lncRNAs (mlncRNAs) and to investigate the mechanism of how the mlncRNAs modulate and switch the functions in the metabolic processes, which is of vital importance for cancer therapeutics and will provide tremendous opportunities for anti-cancer purposes (Du et al, 2013; Liu et al, 2014; Wang et al, 2015; Zhu et al, 2016).

We have demonstrated that using clustering algorithms is able to group proteins into functional modules allowing the identification of MPs (Chapple et al, 2015; Khan et al, 2014; Pritykin et al, 2015). A module corresponds to a functional unit, which is composed of several closely interacted proteins involved in specific tasks in the cell. Therefore, it is promising to use the functional module approach to identify mlncRNAs that exhibit multiple but distinct functionalities. Our study is focused on the moonlighting of human lncRNAs, since lncRNAs are pervasively transcribed in the mammalian genome and several of them play the roles as oncogenic or tumor-suppressor genes in multiple cancers (Ning et al, 2016; Quinn & Chang, 2016; Wahlestedt, 2013; Zhu et al, 2016). We first propose a novel algorithm MoonFinder to identify mlncRNAs that have multiple but unrelated functions. Then, we characterize the sequence features and the localization tendency of these mlncRNA candidates. After that, we construct a mlncRNA-module network and topologically analyzed the mlncRNAs with regard to the association with cancer, diseases, drug targets, and moonlighting proteins. We also predict two cancer lncRNAs exclusively interacted with functional modules from the network.

## 2. Material and methods

### 2.1. Data

#### 2.1.1. Protein-protein interactions

The protein interaction network was constructed using data from InWeb_InBioMap, which is a large-scale, standardized, and transparent resource well suited for functionally interpreting large genomic datasets (Li et al, 2017). It is well known that the protein interaction network is extremely useful for implicating unsuspected pathways in cancer or other disorders, but the currently available datasets all come in different organisms with different interaction number, and the experimental methods are extensive and varied. InWeb_InBioMap contains about 0.6 million interactions between proteins, other than the computationally predicted ones, 57% of them were directly obtained from experiments with human proteins, and 95% from at least one organism, *i.e.*, human, mouse, rat, cow, nematode, fly, or yeast.

#### 2.1.2. Subcellular localization of proteins

The information of protein localization was obtained from the Cell Atlas (Thul et al, 2017), a comprehensive resource for human protein subcellular localization, which is also a subproject of the Human Protein Atlas (Ponten et al, 2008). All proteins were annotated to 14 major compartments and they can be further subdivided into 33 subcellular locations on a single-cell level based on the cellular substructures. We only used the protein annotation of the major compartments.

#### 2.1.3. Protein module identification

We used ClusterONE to identify functional modules from the co-localized protein interaction network (Cheng et al, 2017; Nepusz et al, 2012). It executes a greedy growth algorithm to detect overlapping clusters by starting from a seed. Each processed cluster is supervised by a cohesiveness score to evaluate its separability. Lastly, the cluster pair with a high overlap score (>0.75) is combined and clusters of a small size (<3) and low density (<0.5) are filtered out.

#### 2.1.4. Gene Ontology and functional similarity

The Gene Ontology (GO) provides the functional annotation of gene products (Gene Ontology, 2015; Mazandu et al, 2017). The GO structure is organized as directed acyclic graphs to annotate gene products with appropriate functional terms from three orthogonal ontologies, Biological Process, Molecular Function, and Cell Component. We only used the terms of Biological Process to evaluate the similarity between protein clusters.

The GO semantic similarity provides the basis for functional comparison of gene products or gene product sets. Five common semantic similarity scores are used to measure the functional similarity between identified protein modules, *i.e.*, Resnik, Lin, Jiang and Conrath, Schlicker, and Wang. Wang is a graph-based measurement while the other four are information content based (Yu et al, 2010). Assessment results had shown that one measure may outperform the others in different scenarios in terms of the correlations with sequence similarity, gene co-expression, or interacted gene pairs. Considering the results from different approaches are variable, only the common moonlighting RNAs identified using all the five measurements are determined for further analysis.

#### 2.1.5. lncRNA-protein interactions

The interactions between lncRNAs and proteins were obtained from the database RAID v2.0 (Yi et al, 2017), which is a high-confidence resource of RNA-protein interactions integrating 18 data sources such as StarBase (Li et al, 2014) and LncRNA2Target (Jiang et al, 2015) as well as curated literature. It covers many types of RNAs, such as lncRNA, circRNA, and miRNA, and the interactions between them and proteins are either experimental or computationally predicted. Here, only 12,008 human lncRNAs with protein targets were considered.

#### 2.1.6. Subcellular localization of lncRNAs and RCI

To establish the subcellular localization of lncRNAs, we downloaded data from LncATLAS (Mas-Ponte et al, 2017) and RNALocate (Zhang et al, 2016). Mas-Ponte *et al.* developed a comprehensive resource of lncRNA localization in human cells named LncATLAS (Mas-Ponte et al, 2017). The localization of a lncRNA is represented by its expression level in the RNA-seq data of 15 human cell lines, which is quantified by fragment per kilobase per million mapped (FPKM). They also introduced a measure, Relative Concentration Index (RCI), a log2 transformed ratio of FPKM between two (sub)compartments, to represent the localization tendency of lncRNAs. RNALocate is a localization specific database with manually curated localization classifications across multiple species including Human (Zhang et al, 2016).

#### 2.1.7. lncRNAs biotypes

LncRNAs are usually categorized into six biotypes based on their sequence features such as transcriptional directionality and exosome sensitivity according to FANTOM (Hon et al, 2017). The data were downloaded for assessing whether the lncRNAs of interest are overrepresented in any of these biotypes, *i.e.*, antisense, divergent, sense intronic, intergenic, exonic, and pseudogenes.

#### 2.1.8. Sequence conservation and expression correlation

We collected a set of evolutionarily conserved lncRNAs with correlated expression from lncRNAtor (Park et al, 2014), which serves as a comprehensive resource for functional investigation of lncRNAs. The evolutionary conservation score was calculated for each lncRNAs from UCSC genome database (Pollard et al, 2010) and the co-expression with genes was calculated for the RNA-Seq datasets.

#### 2.1.9. Cancer lncRNAs and drug target proteins

The lncRNAs involved in cancers were obtained from the Lnc2Cancer database (July 4, 2016) (Ning et al, 2016), which curated and integrated the experimental associations between lncRNAs and cancers from the scientific literature. 55 cancer lncRNAs among 76 human cancers were used in this study. Another resource that manually collected the experimentally supported cancer annotations is LncRNADisease (Chen et al, 2013), which contains not only the associations between lncRNAs and cancers but also other diseases. 115 disease lncRNAs implicated in 222 complex diseases, including cancers, are involved in protein interaction network and they were used to evaluate the performance of moonlighting lncRNA candidates. The pharmaceutical drug-targeted proteins were obtained from DrugBank (version 5.0.9) (Law et al, 2014). All drugs in the database are FDA approved. 1,269 proteins in the co-localized protein interaction network are targeted by either small molecule or antibody-based drugs and they are used for further analysis.

#### 2.1.10. Moonlighting proteins

We obtained the moonlighting proteins (MPs) from MoonProt (Mani et al, 2015). MPs are a class of proteins that have multiple but distinct functions that are not due to gene fusions, multiple RNA splice variants or multiple proteolytic fragments. The moonlighting functions of MPs are often not conserved among protein homologues. All the MPs collected in this database are validated biochemically or biophysically.

### 2.2. The workflow of MoonFinder

Like proteins, moonlighting lncRNAs (mlncRNAs) may perform their distinct functions through different target proteins in different cell compartments. Meanwhile, proteins in the interaction network are usually modeled into a variety of functional modules, the proteins in which are associated with specific tasks in the cell or tend to participate in the same biological processes. Hence, it is promising to identify mlncRNAs from the aspect of whether they are targeting functionally unrelated protein modules. MoonFinder integrates several statistical models based on the RNA and protein interactome as well as the protein functional annotations for the identification of moonlighting macromolecules. The workflow of MoonFinder contains the following six steps,

1. Protein interaction network refinement. Refine the protein interaction network by filtering out protein pairs sharing no identical cell compartments and only the co-localized interactions are considered for further analysis.
2. Protein module identification. Detect protein modules from the co-localized protein interaction network (using ClusterONE by default). Each of the detected modules is highly interconnected and expected to be implicated in a specific cellular process.
3. Functional annotation of modules. Establish the annotation of the modules with GO terms by performing the functional category enrichment using hypergeometric test (HGT). The pairwise association between the modules and the GO terms are constructed.
4. Establish RNA-module interactions. Assess whether the target proteins of an RNA are significantly overrepresented in a module. If yes, we define the RNA functionally regulates the module.
5. Construct similarity matrix of modules. All pairwise functional similarities are pre-calculated using five semantic similarity measurements and eventually a similarity matrix is produced.
6. Moonlighting determination. Calculate the principal components (PCs) and their contribution weights of the similarity matrix of the modules targeted by an RNA using principal component analysis (PCA). An RNA is determined as moonlighting if none of the principal components play a dominant role, such as weight > 0.7.

For instance, we say an RNA is moonlighting if the weights of the top three PCs of the module similarity matrix are 0.4, 0.3, and 0.2, respectively, because three PCs (more than one) are unneglectable. On the other hand, an RNA is multifunctional but not moonlighting if the weight of the first PC is 0.95 and the rest share the negligible weight of 0.05, which reflects dependent functions. The workflow of MoonFinder is explained in more detail in Fig. 1.

**Figure 1.**
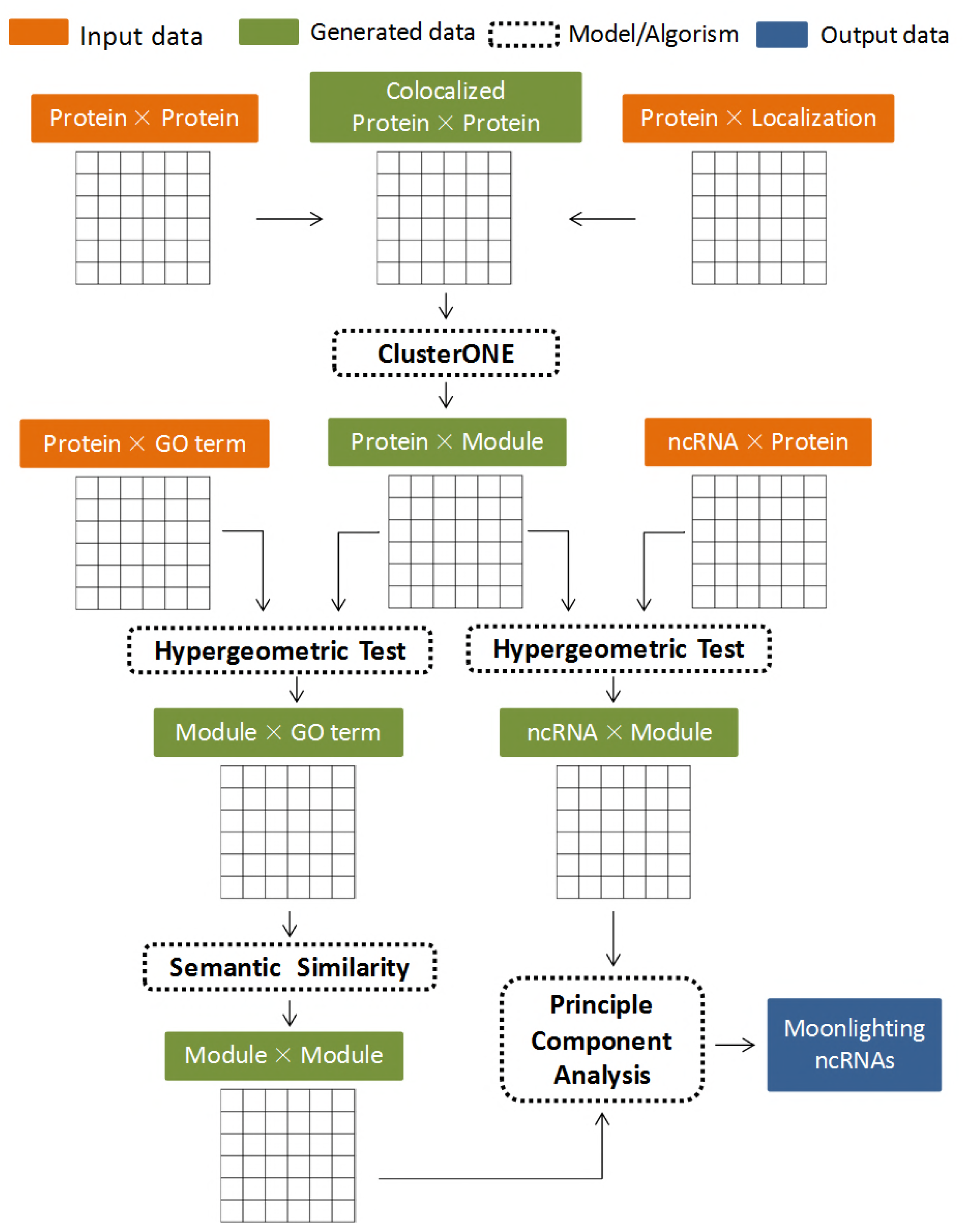
Schematic diagram of MoonFinder. The orange, green and blue boxes represent the input, generated, and output data, respectively. The clustering algorithm and statistical models are shown in the dotted boxes.

### 2.3. Statistical methods

#### 2.3.1. Module-function and lncRNA-module association

The hypergeometric test (HGT) was used to evaluate the statistical significance of the association between lncRNAs and functional modules. A lncRNA is considered to be interacting with a module if the proteins interacting with the lncRNA are significantly enriched in the module, *i.e., P*-value <0.05, which is shown as follows,

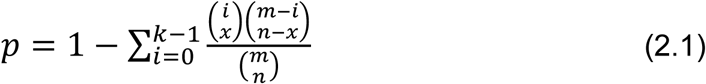

where *n* is the total number of all proteins in the protein interaction network, *m* is the number of proteins analyzed in a module, *x* is the number of proteins interacting with a specific lncRNA, and *i* is the number of proteins in the module interacting with the lncRNA. The *P*-value describes the probability of randomly select no less than *k* interacted proteins in the module with size *m*. Similarly, the model was also utilized to carry out the functional annotation for the identified modules by functional categories (or GO terms), where in this case *n* is still the total number of gene products in the protein interaction network, but *m* represents the module size and *x* represents the term size. *i* is the number of gene products annotated in the term and involved in the module. The *P*-value shows the probability of randomly choosing no less than *k* proteins annotated by a GO term for a given module.

Monte Carlo simulation was adopted to model the probability of interacting mlncRNAs and MPs. *N* pairs of mlncRNAs and MPs were randomly extracted from the lncRNA-protein interaction network and then we calculated the interacting pair number, *T_i_* The *P*-value is the ratio of the number of simulated interactions (*T_i_*) that is larger than the number of practical interactions (*T*), which is mathematically defined as:

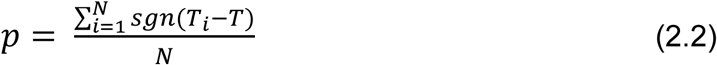

where *sgn* is defined as 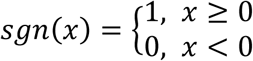

To gain the statistical significance, all comparisons in this study between two lists of genes or gene products were analyzed using Wilcoxon Ranksum Test (WRT, R-3.4.1).

#### 2.3.2. Decomposition of the functional similarity matrix

Principal component analysis (PCA) is a statistical procedure used to reduce the number of features used to represent data. We accomplish the reduction by projecting data from a higher dimension to a lower dimensional manifold such that the error incurred by reconstructing the data in the higher dimension is minimized (Ma & Dai,2011; Ma & Kosorok, 2009). Mathematically, we want to map the features *x* ∈ *R^p^* to 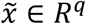 where *q* < *p*. Here, eigenvalue decomposition of the similarity matrix was used to calculate the principle components (PCs) and their weights for the modules interacted by each lncRNA. According to the Spectral Theorem,

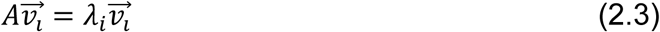

we can calculate the eigenvalues λ_1_+…+λ_m_ of the Similarity matrix (arranged in decreasing order) and accordingly the trace of the matrix is

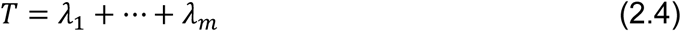

Only the lncRNAs whose similarity matrix can be decomposed into PCs without a dominant role were selected as the candidates, such as the sum of the weights of the first *k* PCs is less than a threshold *τ* (*τ* = 0.7 by default) as follows,

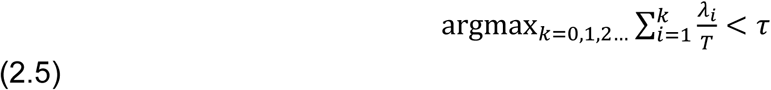

Then the maximum *k* is the number of latent features. The eigenvalue or the weight of a latent feature has minor influence when it is less than 0.1, so we also define λ_*i*_ ≥ 0.1 in Eqt 2.4 and accordingly the number of latent features with key contribution is

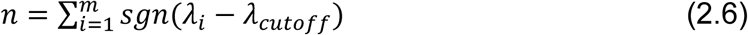

where *sgn* is defined as 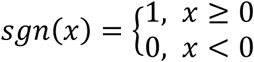 and λ_*_cutoff_*_ = 0.1 number of latent functions for an RNA is

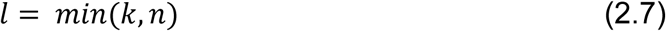

An RNA is determined as moonlighting RNA if *l* is larger than one. Namely, the RNA has more than one latent feature in terms of the interacting functional modules.

#### 2.3.3. Interactor share ratio

To measure how likely the interactors of a given lncRNA are shared by the other lncRNAs, we introduced a score Interactor Share Rate (ISR) as follows,

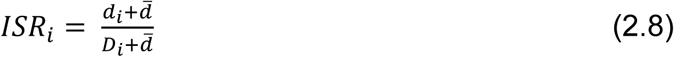

where *d_i_* is the connection degree of RNA *i*, *D_i_* is the sum of the degrees of all neighbors of RNA *i*, and 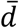 is the average degree of all the RNAs in the network. To make the distribution of the ISR scores more normal and range in between 0 and 1, the scores are normalized as follows,

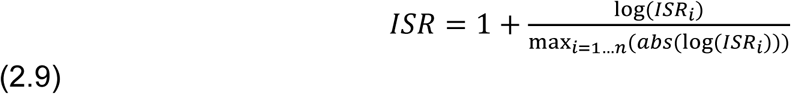

1 means all the neighbors of a lncRNA are only connected to it while a small value, say 0.05, means its neighbors are also connected by many other lncRNAs.

## 3. Results

### 3.1. Experimental and parametric setups of MoonFinder

Moonlighting lncRNAs (mlncRNAs) are assumed to execute multiple distinct functions through interactions with proteins that are localized in different cell compartments, so we identify mlncRNAs from the aspect of whether they are targeting multiple but functionally unrelated protein modules in a co-localized protein interactome. The human protein interactome was first refined and only the co-localized interactions were maintained for module identification, since proteins closely interacted with each other in a module are more likely to reside in the same cell compartment. Eventually, a compartment-specific protein interaction network with 210,410 interactions among 10,111 proteins was constructed. After that, a total of 765 functional modules were identified using ClusterONE, an algorithm that can detect overlapping clusters of proteins highly connected inside but sparse outside (Nepusz et al, 2012). We choose ClusterONE because not only it can detect biological relevant clusters that can be appropriately mapped to modules, but also its ability to softly identify the overlapping modules considering the network topological structure.

Functional enrichment analysis was employed to annotate each identified module with specific and significant function categories. We applied the hypergeometric test (HGT) to obtain the enrichment *P*-values and the FDR adjusted *P*-values of 0.01 were eventually used as the threshold. To establish the RNA-module interactions, similarly, we used HGT to assess whether there are significant connections between the lncRNAs and the functional modules. The target proteins of a lncRNA are significantly overrepresented in a module if the FDR adjusted enrichment *P*-values were less than a given threshold of 0.01. Accordingly, we obtained a bipartite network with 2,726 interactions among 538 lncRNA and 106 protein modules. The function similarity matrices were calculated using the semantic similarities among the modules of each lncRNA using five semantic similarity measurements, *i.e.*, Resnik, Lin, Jiang and Conrath, Schlicker, and Wang (see Section 2.1.4). Only the intersection of the five sets of mlncRNAs identified using the five measures was determined as the candidates (in total 155), because one measure may outperform the others in different expression or interaction scenarios. Importantly, we utilized eigenvalue decomposition to calculate the number of latent features of each lncRNA. The lncRNAs whose module similarity matrix can be decomposed into the principal components without a dominant role (more than one latent feature) were selected as the candidates. The workflow of MoonFinder is described in more detail in the Methods section 2.2 as well as in Fig. 1.

### 3.2. Overview of the mlncRNA candidates

As shown in the Venn diagram of Fig. 2A, among the 1,284 lncRNAs with interacted proteins (background lncRNAs), 538 lncRNAs (flncRNAs) are annotated to at least one functional module and eventually 155 out of them are determined as moonlighting lncRNAs (mlncRNAs), whose target modules are functionally unrelated. These identified mlncRNA candidates are displayed in Supplementary Table 1, including the respective genome locations, cell localization, interacting proteins, and function information. The mlncRNAs were identified using five semantic similarity (SS) measures to quantify the functional similarity between modules. To obtain more reliable results, only lncRNAs in the intersection of the sets of identified mlncRNAs using the five different measures were defined as mlncRNAs (Fig. 2B). The modules targeted by an identical lncRNA have a higher probability of sharing the same functions than the randomly picked modules. Not surprisingly, significantly more mlncRNAs are detected when simulating the randomly selected modules as the target modules. Specifically, Fig. 2C shows that around 40% of the lncRNAs with target module can be identified as mlncRNAs using either SS measures, the ratio is as low as 28% when using Lin’s measure, while the ratios increase to about 60% when the target modules are randomly picked.

**Figure 2.**
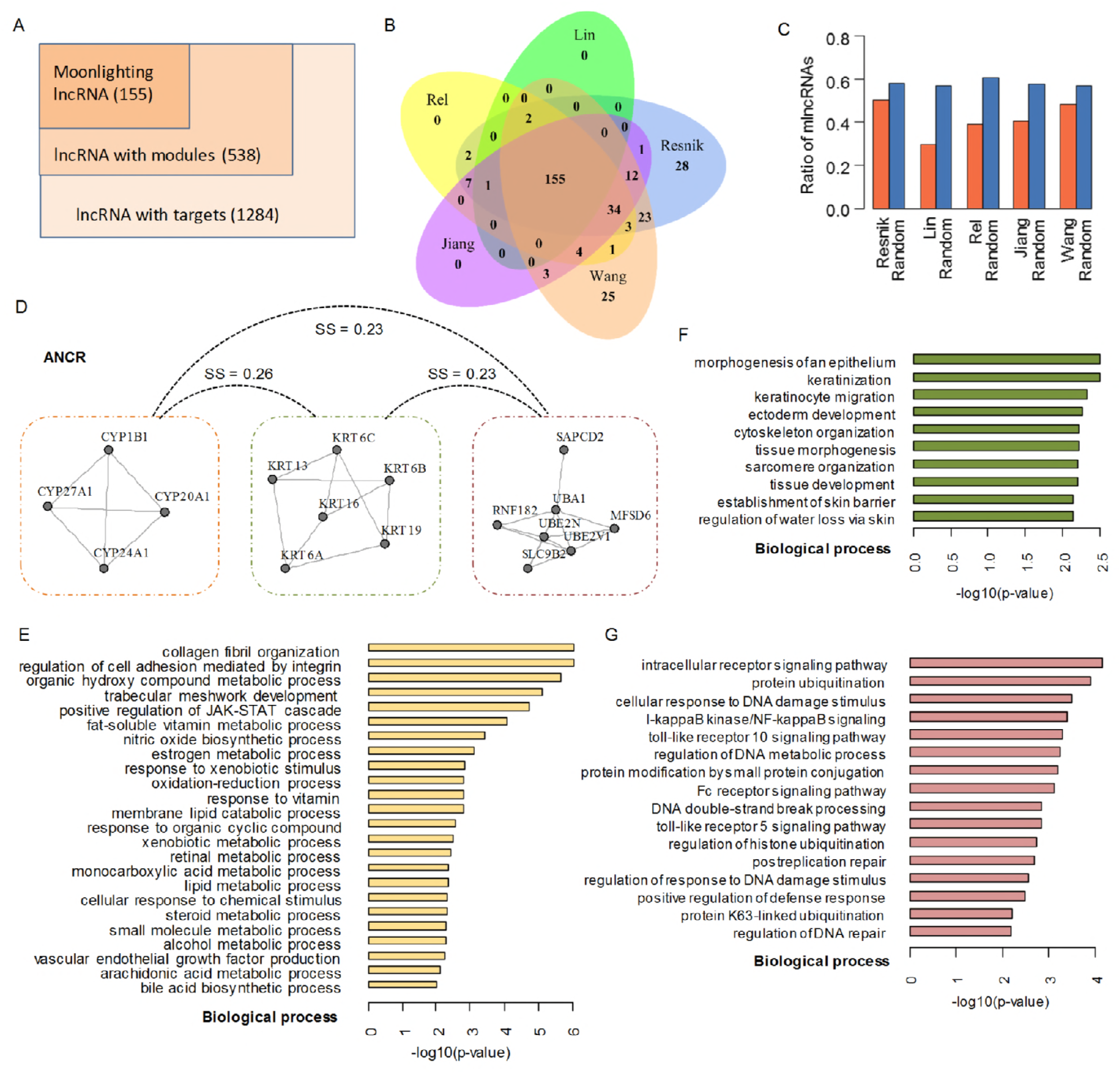
Overview of the identified mlncRNAs. (**A**) Venn diagram of the lncRNAs. (**B**) Venn diagram of the mlncRNAs identified using five semantic similarity measures. (**C**) The ratio of real and randomly identified mlncRNAs. (**D**) An example of mlncRNA *ANCR*. (**E-G**) Gene Ontology functional enrichment of the three modules regulated by *ANCR*.

Here, we take *ANCR* as an example to illustrate its moonlighting functions. *ANCR* is an anti-differentiation lncRNA that is required to enforce the undifferentiated state in somatic progenitor populations of epidermis. Using MoonFinder, we observed *ANCR* mainly interacts with three functional modules with closely linked proteins inside and no proteins are shared between any two modules, as shown in Fig. 2D. The SS scores are extremely low among the three modules, *i.e.*, 23%, 26%, and 23%, respectively, owing to few GO terms of biological process are hierarchically correlated (Fig. 2, E-G). Specifically, one module was enriched in a variety of metabolic processes, such as estrogen, retinal, and steroid, *etc.*, whereas another module was enriched in functions like tissue morphogenesis and development. Signaling pathways for enforcing intracellular receptors like toll-like receptor 5 and 10 were highly represented for the other module. Consequently, *ANCR* was determined as a mlncRNA who shows its functional diversity via regulating protein modules taking part in distinct biological processes and it would serve as a highly reliable candidate of moonlighting lncRNA.

### 3.3. Sequence features of mlncRNAs

To investigate whether the mlncRNAs form a distinct group of lncRNAs, we analyzed the sequences of the corresponding non-coding genes to detect common features such as biotype, gene length, transcript length, transcript number, exon length, and exon number, as well as the evolutionary conservation and expression correlation with orthologous genomes. The candidates were compared with other two groups of lncRNAs, *i.e.*, the entire set of lncRNAs with protein targets (background lncRNAs) and the functional module related lncRNAs (flncNRAs), which interacts with functionally related or unrelated modules (Fig. 2A). Hence, it is important to identify the unique characteristics of mlncRNAs that the other types of lncRNAs do not exhibit. Although no significant differences of gene length were detected for distinct categories of lncRNAs (Fig. 3A), the candidate mlncRNAs have a significantly longer transcript than the other types of lncRNAs. On average, the transcript length is about 19,500 compared with about 11,000 for the flncRNAs (WRT *P*-value = 1.27e-3; Fig. 3B). But the number of transcripts in these lncRNA categories are similar to each other, only a marginal significance was tested between the mlncRNAs and the background lncRNAs (WRT *P*-value = 4.7e-2; Fig. 3C). Importantly, we observed that the exon of mlncRNAs are significantly shorter than the other categories of lncRNAs (WRT *P*-value = 3.4e-6 and *P*-value = 8.3e-9; Fig. 3D) but the number is much more than the regular lncRNAs (10 vs 1, WRT *P*-value = 2.2-e16; Fig. 3E), probably owing to the transcripts of mlncRNAs are on average much longer than that of the regular lncRNAs. In short, mlncRNAs have short but more exons, which is a potential sequence feature for lncRNAs to moonlight in between multiple biological functions.

**Figure 3.**
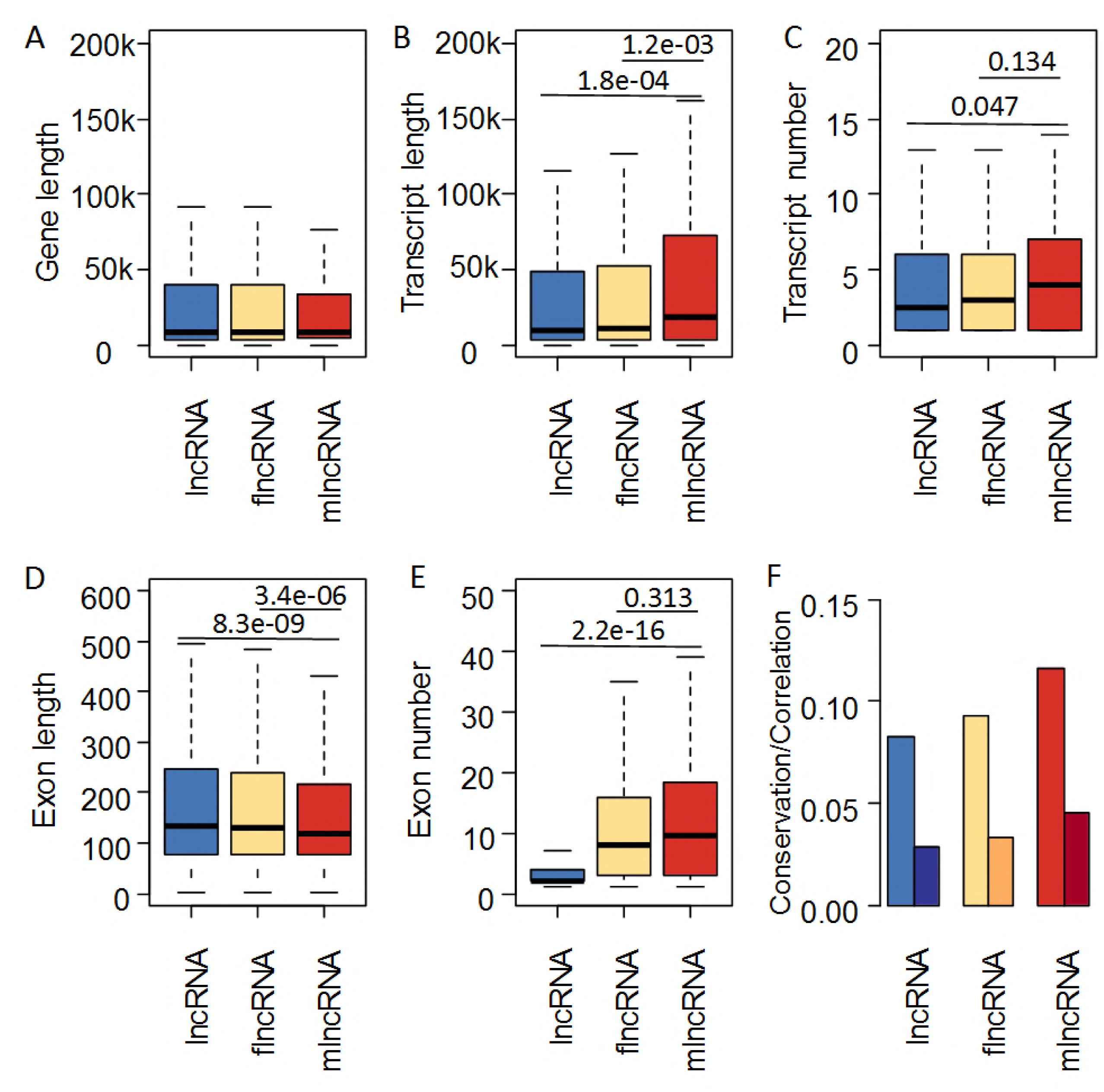
The distributions of lncRNAs in different functional groups with regard to distinct sequence features. (**A**) Gene length. (**B**) Transcript length. (**C**) Transcript number. (**D**) Exon length. (**E**) Exon number. (**F**) Evolutionary conservation and expression correlation. Outliers are not shown. Blue, yellow, and red boxes (or bars) represent lncRNA, flncRNA, and mlncRNA, respectively.

Phylogenetic conservation and expression correlation are strong evidence for inferring functions of coding or non-coding genes (Cheng et al, 2016a; Cheng et al, 2016b; Park et al, 2014). As lncRNAs are crucial in biological processes if they are evolutionarily conserved or expression-correlated across species, we checked whether the mlncRNAs tend to be conserved among orthologous genomes and whose expression patterns are highly correlated in orthologs. As shown in Fig. 3F, the mlncRNAs are prone to gain higher conservation scores (>0.12) than flncRNAs and background lncRNAs (<0.10). The scores of flncRNAs are lower than mlncRNAs but still higher than that of the background lncRNAs. Similarly, the largest proportion of mlncRNAs are found to be expression conserved and the ratio of flncRNAs is second to it. Consequently, the evolution and expression pattern of mlncRNAs is more conserved than the other lncRNAs, which is in contrast with the conventional knowledge that lncRNAs are generally less conserved than mRNAs and proteins (Hon et al, 2017; Park et al, 2014), revealing that the lncRNAs moonlighting in the cells may play more important biological roles. Besides, RNA species were officially grouped into several biotypes by their transcriptional direction and exosome sensitivity (Hon et al, 2017). Here we also examined the relationship between the functional categories and biotypes, but no significant correlation was detected (Supplementary Fig. 1A).

### 3.4. Subcellular localization features

Next, we aimed to understand how the mlncRNAs behave relative to the other lncRNAs in terms of subcellular localization. To investigate the spatial distribution of lncRNAs at a subcellular level, we applied Relative Concentration Index (RCI) (Mas-Ponte et al, 2017), a ratio of a transcript’s concentration between two cellular compartments, to measure the localization tendency of non-coding RNAs. Essentially, RCI is the log2 transformed ratio of FPKM (fragments per kilobase per million mapped) in two compartments like cytoplasm and nucleus. First, we calculated the cytoplasmic-nuclear RCI to measure the relative concentration of a lncRNA between the cytoplasm and the nucleus in 15 cell lines. Fig. 4A illustrates the RCI distributions of mlncRNAs, flncRNAs, and background lncRNAs. It is apparent that mlncRNAs tend to have higher RCI values compared to the other two categories of lncRNAs in almost all these cell lines except SK.N.DZ, SK.MEL.5, and K562, indicating that the mlncRNAs are more likely to reside in the cytoplasmic in comparison with the other lncRNAs. Then, we further investigated the localization of mlncRNAs at the sub-compartment level, since LncATLAS also provides information about enrichment in the cytoplasmic and nuclear sub-compartments of the K562 cells. As shown in Fig. 4B, the sub-nuclear RCI values of mlncRNAs are higher than that of the other two groups of lncRNAs while the sub-cytoplasmic RCI values are relatively small. Namely, in the nucleus, the mlncRNAs are prone to appear in the sub-compartments of nucleoplasm, nucleolus, and chromatin, whereas in the cytoplasm, the mlncRNAs are not likely to reside in insoluble and membrane relative to the other lncRNAs. That is why the cytoplasmic-nuclear RCI values of mlncRNAs are almost the same as the background lncRNAs in the K562 cell line (Fig. 4A). Meanwhile, we also calculated the expression value distribution of lncRNAs in each sub-compartment in the K562 cell line. More importantly, the expression values of the mlncRNAs are significantly higher in all the sub-compartments, revealing that the expression abundance of lncRNAs is crucial in executing the part-time functions (Fig. 4C).

**Figure 4.**
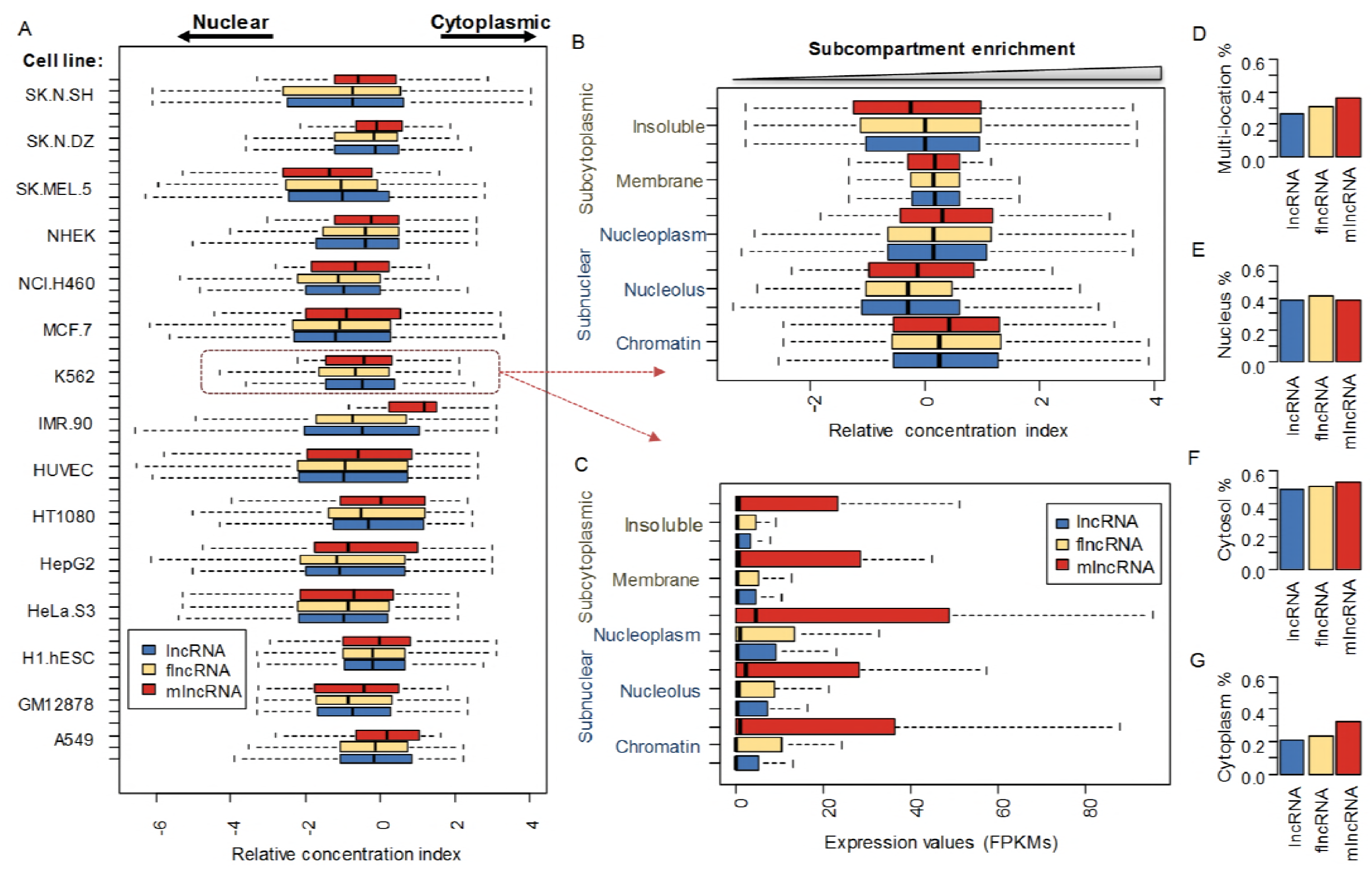
The mlncRNA localization and expression features. (**A**) RCI distribution of all lncRNAs (blue), flncRNAs (yellow) and mlncRNAs (red) for each cell line. (**B**) Sub-compartment expression value distribution of lncRNAs in the K562 cell line. (**C**) Sub-compartment RCI distribution of lncRNAs in the K562 cell line. (**D**) The ratios of different groups of lncRNAs residing in multiple locations. (**E-G**) The ratios of different groups of lncRNAs residing in the nucleus, the cytosol, and the cytoplasm, respectively. RCI, Relative Concentration Index

In addition, we also used another RNA localization resource RNALocate, which contains manually-curated localizations classifications, to investigate the localization tendency of mlncRNAs. In this database, the lncRNAs were collected and annotated to different cell compartments, e.g., nucleus, cytosol, and cytoplasm. We calculated the ratio of lncRNAs in these compartments separately for each category of lncRNAs. We found that mlncRNAs tend to appear in more than one compartment and localize in the cytoplasmic compartments such as cytosol and cytoplasm (Fig. 4D-G). The ratio of multilocation mlncRNAs is as high as 0.35, which is much higher than that of flncRNAs and the background lncRNAs (about 0.3 and 0.26, respectively; Fig. 4D). More importantly, mlncRNAs were found to be enriched in cytosol and cytoplasm with the ratios of 0.55 and 0.3 (Fig. 4F, G), respectively, whereas the ratio is comparable to the other categories of lncRNAs in the nucleus (Fig. 4E). Consequently, we can draw the same conclusion that mlncRNAs have a localization tendency of residing in the cytoplasmic compartment.

### 3.5. Topological features of mlncRNAs and its roles in cancers

lncRNA functions through its interacting partners. Accumulating studies show that the multi-functionality of lncRNAs as interacting hubs with other molecules such as proteins, DNAs, and RNAs. Apparently, the candidate mlncRNAs connect a significantly larger number of proteins and modules than the other lncRNAs according to the identification methodology. On average, the number of partner proteins is 36.1 for mlncRNA while less than 20 for the others (WRT *P*-value = 7.4e-8; Supplementary Fig. 1B). The number of the interacted module is around 6.8 for mlncRNA compared with 5 for the other lncRNAs (WRT *P*-value = 8.8e-14; Supplementary Fig. 1C). To illustrate the combinatorial regulation and give a systematic description, we constructed a mlncRNA-module regulatory network, in which the edges link the candidate mlncRNAs to their corresponding functional modules. This network contains in total 1,055 predicted regulatory interactions between 155 mlncRNAs and 83 modules (Fig. 5A). Some modules connect more candidate mlncRNAs than others, indicating that they might be engaged in a larger number of moonlighting regulations. In particular, the largest module in the center of the network shows the highest degree of 70, suggesting that it is subjected to the regulations of 70 mlncRNAs. The proteins in this module were found to be mainly implicated in biological processes such as nucleic acid metabolic process, gene expression, and amniotic stem cell differentiation (Fig. 5B).

**Figure 5.**
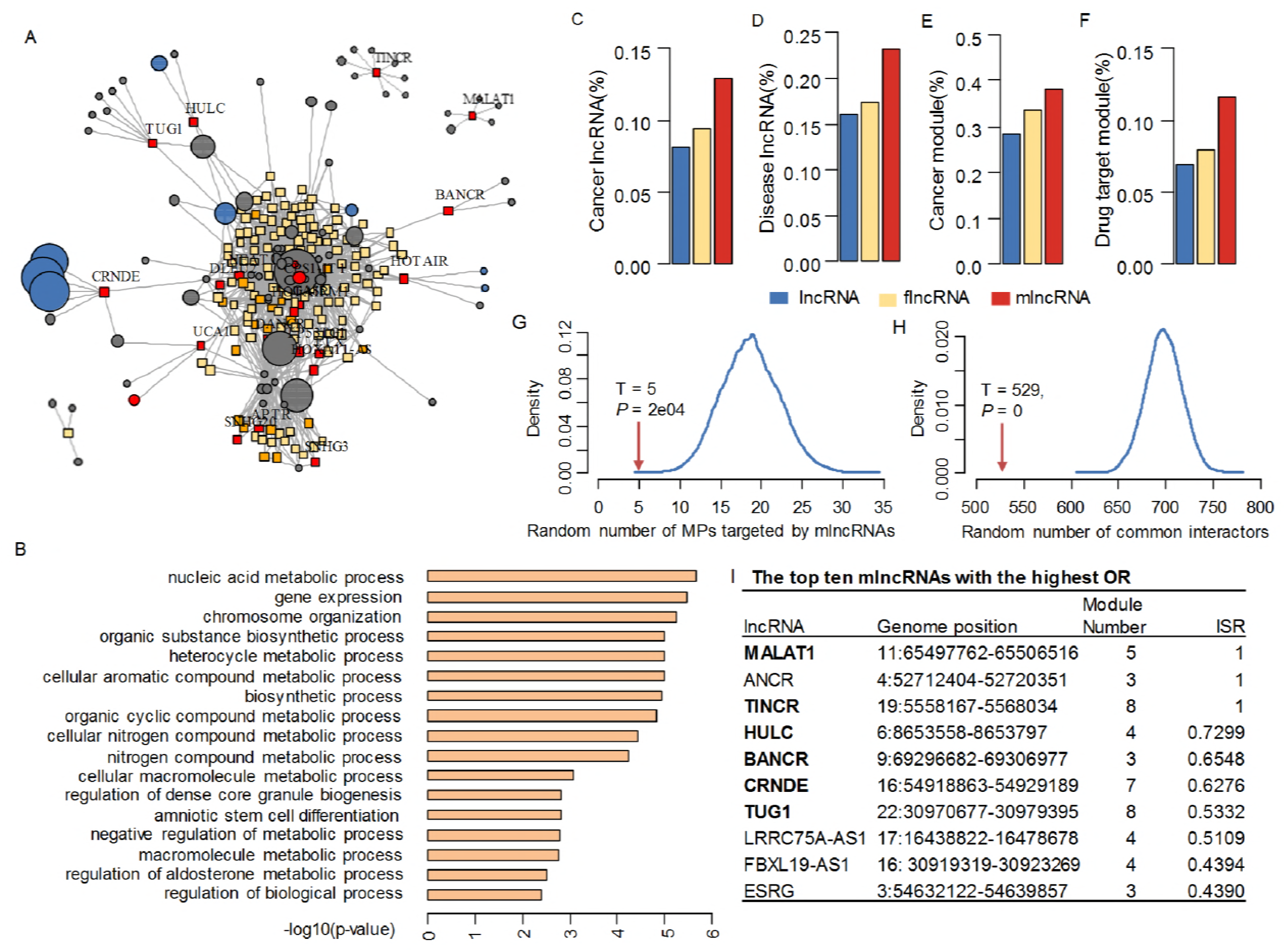
Association between mlncRNAs and diseases. (**A**) The mlncRNA-module regulation network. RNAs are represented in squares while modules are in circles. RNAs associated with cancer and diseases are shown in red and orange, respectively. The circle size corresponds to the module size. (**B**) Gene Ontology function enrichment of the module with the largest number of regulated mlncRNAs. (**C, D**) The ratio of cancer and disease lncRNAs among the three lncRNA categories. (**E, F**) The ratio of lncRNAs associated with cancer and drug target module. (**G**) mlncRNAs tend not to interact with MPs. T, the number of practical mlncRNA-MP interactions. (**H**) mlncRNAs and MPs tend to share less interacting partners. T, the number of common partners practically interacted by mlncRNAs and MPs. (**I**) Summary of the top ten mlncRNAs with the highest OR scores.

To determine whether the moonlighting of lncRNAs is implicated in the formation and development of cancers and other diseases, we associated the mlncRNAs with public available cancer and disease lncRNAs as well as cancer proteins and drug target proteins (see Methods). Around 13% of the mlncRNA candidates have been studied involved in cancer processes, while the ratios of flncRNAs and background lncRNAs are less than 10% (Fig. 5C). When considering other complex diseases, not surprisingly, the ratio is as high as 23% for mlncRNAs, which is still much higher than that the other categories (16% and 17%, respectively; Fig. 5D). From the perspective of regulated functional modules, about 39% of the candidates are included in modules significantly enriching cancer proteins, whereas the ratios decrease to about 29% and 33% for the other two groups (Fig. 5E). Similarly, for the modules enriched of drug targets, they are more likely to interact with mlncRNAs than the flncRNAs and the background lncRNAs (12% vs 7% vs 8%; Fig. 5F). Besides, we can draw the same conclusion when concentrating on the proportion of cancer modules or drug target modules regulated by the mlncRNAs (Supplementary Fig. 1D and E). Therefore, these results demonstrate that the mlncRNAs exercise a great influence on cancer metastasis and progression through pairwise interactions and clustered modules of proteins in the regulatory network.

More importantly, we observed that the mlncRNAs and moonlighting proteins (MPs) tend to be mutually exclusive in terms of interactions and interacted partners. Only five MPs directly interacts with the mlncRNAs, whereas on average the value is as high as 18.9 when randomly selected the same number of MPs (Monte Carlo *P*-value = 1e-04, HGT *P*-value=5.4e-18, Fig. 5G). Also, we simulated the interacted partners of both mlncRNAs and MPs 10000 times and found the number of common partners between them is 691 on average, which is significantly higher than the practical value of 529 (Monte Carlo *P*-value = 0, HGT *P*-value=2.1e-86, Fig. 5H). In other words, the number of common partner proteins that the moonlighting lncRNAs and proteins shared is significantly less than that of randomly selected ones. These results indicate the mechanism that the cells make full use of the macromolecules to efficiently and systematically perform cellular tasks avoiding the redundant implementations.

Additionally, from the mlncRNA-module network in Fig. 5A, we found the mlncRNAs that exclusively interact functional modules tend to be cancer-related. Accordingly, we introduced a score, Interactor Share Rate (ISR), to measure how likely the interactors of a given lncRNA are shared by the other lncRNAs (see section 2.3.3). We found that the cancer mlncRNAs have significantly higher ISRs than that of the others (WRT *P*-value=3.2e-03, Fig. S1F). For the mlncRNAs with the top ten highest ISR scores, six out of them are cancer lncRNAs (Fig. 5I). When strengthening the threshold to 0.5, six out of eight (75%) of the mlncRNAs are cancer genes and the other two, *ANCR* and *LRRC75A-AS1*, could be considered as the candidates of cancer mlncRNA, where the dysfunction or inappropriate switching of these RNAs in different cell compartments may result in the biological activity of cancer, although further experimental works are needed to warrant this claim.

## 4. Discussion

In this study, we introduced a computational framework MoonFinder to systematically identify moonlighting lncRNAs (mlncRNAs) based on the integrated lncRNA and protein interaction network as well as the protein functional annotations. In total 155 lncRNAs were determined as candidates with multiple but distinct functions. Also, we characterized them from various aspects of sequence features, evolutionary conservation, expression correlation, expression abundance, localization tendency, and interaction patterns, which will facilitate our further understanding of their functions and the mechanism of moonlighting.

Remarkably, we observed that the non-coding genes that transcript mlncRNAs tend to have shorter but more exons, which is a potential sequence feature for lncRNAs to moonlight in between multiple biological functions. Also, we found the evolution and expression patterns of mlncRNAs are more conserved than the other lncRNAs, which in contrast with the conventional knowledge that lncRNAs are generally less conserved than mRNAs and proteins(Hon et al, 2017; Park et al, 2014), suggesting that mlncRNAs are central for the homeostasis maintenance of human.

More importantly, we found that mlncRNAs have a localization tendency of residing in cytoplasmic compartment, although they display high expression across all the cell compartments. mlncRNAs are expressed significantly higher in all the sub-compartments of the K562 cell lines in comparison with the other lncRNAs, suggesting that the high expression abundance is necessary for executing the part-time functions. We studied the localization tendency and translocation activity of these mlncRNAs because lncRNAs are diversely resided in the cells and play a crucial role as modulators to regulate gene expression in multiple ways (Cabili et al, 2015; Ferre et al, 2016; Quinn & Chang, 2016; Zhou et al, 2017b; Zhu et al, 2016). lncRNAs have a variety of subcellular localization patterns, which are not limited to specific nuclear and cytoplasm localization but also nonspecific localization in both the nucleus and cytoplasm (Barabasi & Oltvai, 2004; Buxbaum et al, 2015). For the lncRNAs localized in multiple compartments, in the future we will investigate whether the intercommunication can modulate the interaction pattern or expression abundance, e.g. regulating the abundance of lncRNAs in one compartment may influence the function of the other cell compartment.

Our result also shows that mlncRNAs and MPs are rather mutually exclusive in terms of their direct interactions and interacting partners. In other words, lncRNAs and proteins with moonlighting functions are not likely to interact with each other and they even tend to share fewer neighbors in the regulatory network. The reason might be that the macromolecules in cells are usually organized to be efficient to perform different cellular tasks without redundancy. According to the mlncRNA-module bipartite network, we also predicted eight cancer lncRNAs and six out of them were previously identified as cancer lncRNAs by different experimental assays.

We believe our observations can aid our and other research groups to understand how they function in a moonlighting manner and help in designing RNAs with novel functions and applications. Moreover, investigating the mechanisms that determine the functional diversity of mlncRNAs has the potential to provide new insights into their biogenesis, physical interaction, subcellular localization, and therapeutic application in diseases. In the future, we will investigate the mechanism of how the mlncRNAs modulate and switch the functions in metabolic processes, which is of vital importance for cancer therapeutics and will provide tremendous opportunities for anti-cancer strategies. The moonlighting feature of the other types of RNAs, such as miRNA and circRNA (Chen, 2016a), will also be studied and compared and eventually a moonlighting atlas of both RNAs and proteins will be provided.

## Acknowledgements

This work was supported by The Chinese University of Hong Kong Direct Grant; and the Research Grants Council of Hong Kong GRF Grant [414413].

## Author contributions

L.C. and K.L. conceived and designed the experiments. L.C. analyzed the data, performed the experiments, and analyzed the results. K.L. provided the project supervision. L.C. and K.L. wrote the manuscript. All authors reviewed and approved the final manuscript.

Competing financial interests: nothing declared.

